# Rassf1a is required for the maintenance of nuclear actin levels

**DOI:** 10.1101/559310

**Authors:** Maria Chatzifrangkeskou, Dafni-Eleftheria Pefani, Michael Eyres, Iolanda Vendrell, Roman Fischer, Daniela Pankova, Eric O’Neill

## Abstract

Nuclear actin participates in many essential cellular processes including gene transcription, chromatic remodelling and mRNA processing. Actin shuttles into and out the nucleus through the action of dedicated transport receptors importin-9 and exportin-6, but how this transport is regulated remains unclear. Here we show that RASSF1A is a novel regulator of actin nucleocytoplasmic trafficking and is required for the active maintenance of nuclear actin levels through supporting binding of exportin-6 (XPO6) to RAN GTPase. *RASSF1A* (Ras association domain family 1 isoform A) is a tumour suppressor gene frequently silenced by promoter hypermethylation in all major solid cancers. Specifically, we demonstrate that endogenous RASSF1A localises to the nuclear envelope (NE) and is required for nucleo-cytoplasmic actin transport and the concomitant regulation of Myocardin-related transcription factor A (MRTF-A), a coactivator of the transcription factor serum response factor (SRF). The RASSF1A/RAN/XPO6/nuclear actin pathway is aberrant in cancer cells where *RASSF1A* expression is lost and correlates with reduced MTRF/SRF activity leading to cell adhesion defects. Taken together, we have identified a previously unknown mechanism by which the nuclear actin pool is regulated and uncovered a previously unknown link of RASSF1A and MTRF/SRF in tumour suppression.

## Introduction

Actin is one of the most highly conserved cytoskeletal proteins and found in all eukaryotic cells. The fundamental roles of actin are critical for biological processes such as determination of cell shape, vesicle trafficking and cell migration-processes often deregulated in transformed cells. While the role of cytoplasmic actin is well established, the presence of nuclear actin is increasingly appreciated to play a crucial role in cellular responses to internal and external mechanical force. Nuclear actin is actively imported and exported through the activity of importin-9 (Dopie *et al*, 2012) and exportin-6 (Stuven *et al*, 2003) and has been implicated in transcription (Egly *et al*, 1984; Hofmann *et al*, 2004), cellular differentiation (Sen *et al*, 2015) and DNA repair (Belin *et al*, 2015). Nuclear actin monomers have been shown to directly regulate the Myocardin-related transcription factor A (MRTF-A), a mechanosensitive co-factor of the serum response factor (SRF) transcription pathway (Vartiainen *et al*, 2007b; Baarlink *et al*, 2013).

The Ras-association domain family (RASSF) genes are upstream regulators of the Hippo tumour suppressor pathway. The family is composed of ten members which are divided into two subgroups: i) RASSF 1–6 possess a Ras Association (RA) domain and a SARAH (<u>Sa</u>v/<u>Ra</u>ssf/<u>H</u>po) protein-protein interaction domain in the C-terminus, ii) RASSF 7–10 also contain a RA domain at the N-terminus but lack a recognisable SARAH domain (Sherwood *et al*, 2010). The RA domains of the RASSF family bind K-RAS, H-RAS, RAP1/2 and RAN GTPases with varying affinity (Avruch *et al*, 2006; Dallol *et al*, 2009). The RASSF1A isoform is a *bone fide* tumour suppressor gene whose inactivation is implicated in the development of a wide range of human tumours including breast, lung and gastrointestinal cancer (Grawenda & O’Neill, 2015). Although gene deletion and germline mutations exist, the most widespread loss of RASSF1A function occurs through promoter hypermethylation associated transcriptional silencing (Grawenda & O’Neill, 2015). RASSF1A directly binds Hippo kinases, mammalian sterile 20-like kinase 1 and 2 (MST1 and MST2) through the SARAH domain, promoting downstream hippo pathway signaling to YAP1 (Guo *et al*, 2007; Matallanas *et al*, 2007). In response to DNA damage, nuclear RASSF1A is phosphorylated on Ser131 by ATM (ataxia-telangiectasia mutated) or ATR (ATM- and Rad3-Related) kinases, which promotes dimerization and trans-autophosphorylation of MST2 required for its activation (Pefani *et al*, 2014; Hamilton *et al*, 2009). Plasma membrane localisation of cytoplasmic RASSF1A occurs in response to growth factor signaling or direct RAS association with the RA domain (Pefani *et al*, 2016). However, the localisation of RASSF1A to mitotic structures has been attributed to RA domain binding to tubulin (also a GTP binding protein) and is a likely cause of hyperstabilization of tubulin commonly observed in interphase cells overexpressing exogenous levels (Liu *et al*, 2003; Dallol *et al*, 2004; Vos *et al*, 2004; El-Kalla *et al*, 2010).

Here we show that nuclear RASSF1A is recruited to the NE by a lamina-associated pool of the hippo kinase, MST2. Furthermore, we find that RASSF1A is required for the association of exportin-6 (XPO6) with RAN-GTPase, the nuclear export of the XPO6 cargoes, actin and profilin, an actin-binding protein involved in the dynamic sensing of the actin cytoskeleton and the regulation of the MRTF/SRF axis. Tumour cells lacking RASSF1A expression display high levels of nuclear actin/profilin, reduced MRTF levels and low transcription of SRF target genes, including SRF itself. In line with these findings, we find that human tumours display a high level of correlation between RASSF1A and SRF expression, with evidence for low SRF mRNA as a poor prognostic factor in breast and liver cancers. These events reveal the potential primary mechanism for the tumour suppressor RASSF1A in cancer being mediated through deregulation of nuclear actin transport.

## Results

### RASSF1A localises at the inner nuclear membrane

RASSF1A acts as a microtubule-associated protein that stabilises microtubules (Dallol *et al*, 2004; Vos *et al*, 2004; El-Kalla *et al*, 2010; Liu *et al*, 2003). However, these phenotypes were determined after the accumulation of overexpressed protein over a 24–48 h period. To identify the localisation of endogenous nuclear RASSF1A by indirect immunofluorescence (IF) and methanol fixation, we used an anti-RASSF1A antibody (ATLAS), the specificity of which was validated in cells depleted for RASSF1A (Figs EV1A and B). Interestingly, in HeLa cells endogenous RASSF1A is distributed predominantly throughout the nuclear interior and at the perinuclear regions. Co-staining of RASSF1A and the inner nuclear membrane (INM) protein Lamin A/C showed a high degree of co-localisation and similar fluorescence intensity profiles (Fig 1A). Silencing of RASSF1A resulted in loss of the nuclear ring staining, indicating that the fluorescence signal is specific for RASSF1A (Fig EV1B). Localisation was conserved across multiple cell types including mouse embryonic fibroblasts (MEFs), human retinal pigmented epithelial RPE-1 cells and human embryonic kidney HEK 293 cells (Fig 1B). The *RASSF1A* gene promoter is highly methylated in the breast carcinoma cell line MDA-MB-231 and therefore significantly downregulated (Montenegro *et al*, 2012). Concomitantly, no immunofluorescence was observed in MDA-MB-231 cells, but upon transient transfection with a plasmid expressing RASSF1A, NE localisation of the exogenous protein was evident (Fig 1C). Alternatively, treatment of MDA-MB-231 with the demethylating agent 5’-aza-dC led to the re-expression of endogenous *RASSF1A* and yielded similar staining (Fig EV1C).

**Figure 1:**
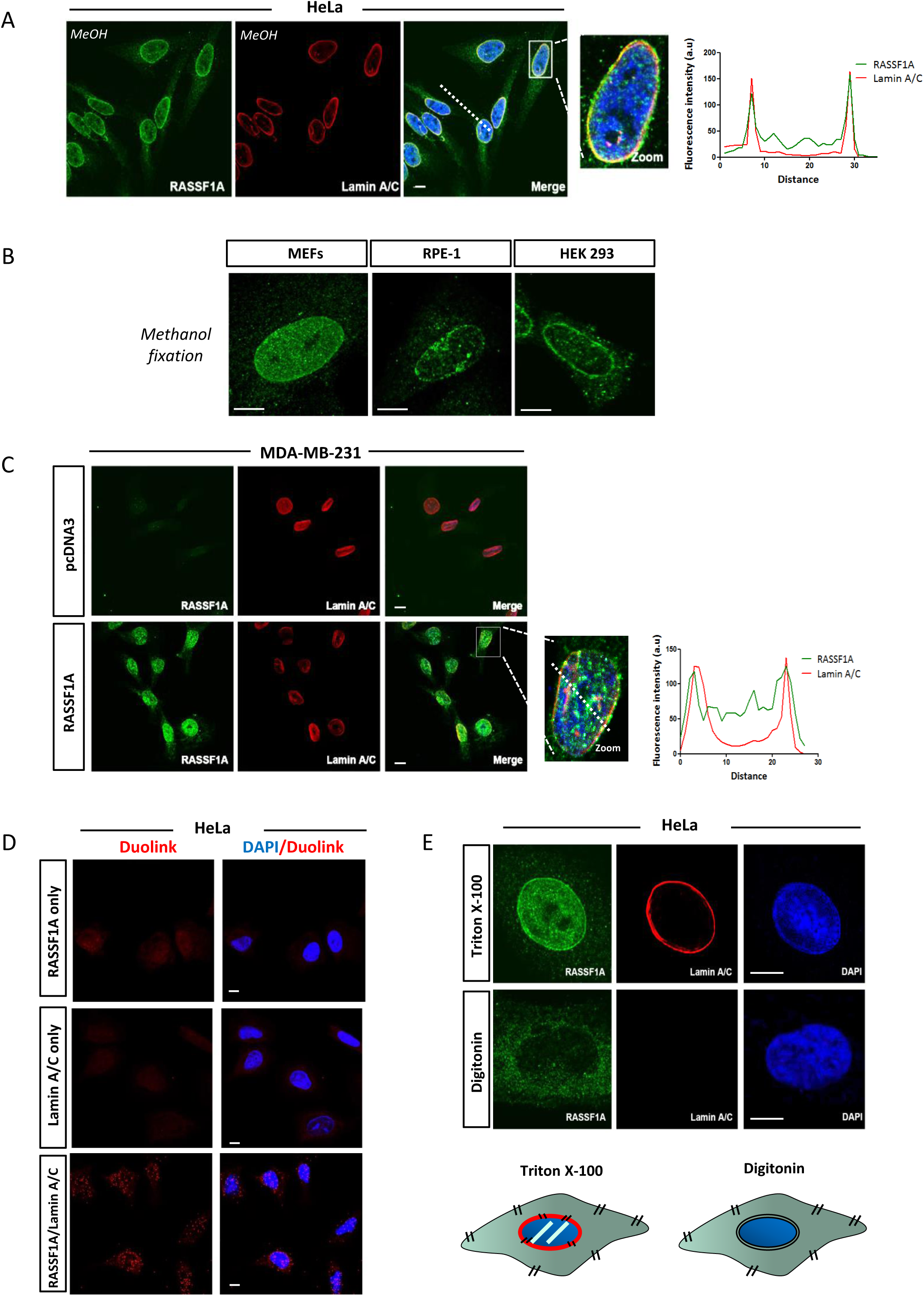
RASSF1A localises at the inner nuclear membrane. A. Representative confocal images of RASSF1A localisation in HeLa cells. Nuclear DNA was stained with DAPI. Fluorescence intensity profile of Lamin A/C (red) and RASSF1A (green) signals across the HeLa nuclei. Position of line scan indicated by the dashed white line. B. Immunofluorescence images of endogenous RASSF1A expression in mouse embryonic fibroblasts (MEFs) (left), human retinal pigmented epithelial RPE-1 cells (middle) and human embryonic kidney HEK293 cells (right). C. Co-staining of endogenous *RASSF1A* expression in MDA-MB-231 cells transfected with the pcDNA3 expressing RASSF1A. Western blot analysis shows the protein levels of RASSF1A following plasmid transfection. Fluorescence intensity profile of Lamin A/C (red) and RASSF1A (green) signals across the HeLa nuclei. Position of line scan indicated by the dashed white line. D. Duolink PLA assay was conducted in HeLa cells with the Duolink II red kit, and cells were imaged in the red fluorescence channel. Immunofluorescence images of the fluorescent signal obtained for single primary antibody controls (RASSF1A only, Lamin A/C only) and interaction between RASSF1A and Lamin A/C (RASSF1A/Lamin A/C) E. *Upper:* Representative confocal images of HeLa cells permeabilised either with Triton X-100 (top panels) or digitonin (bottom panels) and stained for RASSF1A, Lamin A/C and DNA (Dapi). Scale bars = 10 µm. *Lower:* Schematic representation of the cellular membranes permeabilised with Triton X-100 and digitonin.

We next validated the NE association of RASSF1A, by investigating the co-localisation of RASSF1A with Lamin A/C, using proximity ligation assay (PLA) in HeLa cells. We found that RASSF1A is closely localised to Lamin A/C at the nuclear envelope (NE) (Fig 1D), but associates independently as Lamin A/C knockdown did not affect localisation (Fig EV1 B). These results suggest that Lamin A/C is not necessary for proper targeting of RASSF1A to the NE. To further assess the localisation of RASSF1A in relation to the structure of the NE, HeLa cells were treated with either Triton X-100 or digitonin. Whereas Triton X-100 is used to permeabilise all cellular membranes, digitonin can permeabilise preferentially the plasma membrane of cultured cells and leave the NE intact. Therefore, only proteins of the outer nuclear membrane (ONM) that face the cytoplasm are detectable in digitonin-permeabilised cells. In Triton X-100-permeabilised HeLa cells RASSF1A co-localises with Lamin A/C (Fig 1E). In contrast, neither Lamin A/C nor RASSF1A were detected at the nuclear envelope in digitonin-permeabilised cells. These results support the hypothesis that RASSF1A localises at the inner nuclear membrane (INM) of the NE (Fig 1E).

The ATM and ATR kinases catalyse phosphorylation of RASSF1A at Ser131 to regulate its activity (Hamilton *et al*, 2009). Phosphorylated (Ser131) RASSF1A was present within the nucleus and at the INM of HeLa cells (Fig EV1 D). To determine whether phosphorylation of RASSF1A on Ser131 by ATM and ATR kinases is required for its localisation at the inner nuclear membrane, HeLa cells were treated with the ATR inhibitor (VE-821) or ATM inhibitor (KU55933). Inhibition of ATR, ATM or combined ATR/ATM activity did not abolish RASSF1A perinuclear localisation (Fig EV1 E). Collectively, the above observations show that the RASSF1A association with the NE is specific and not dependent on the kinase activity of ATR and ATM kinases.

### RASSF1A localisation at the NE is mediated by MST2

RASSF1A directly binds Mammalian STE20-Like Protein Kinases 1 and 2 (MST1 and MST2) to support maintenance of MST1/2 phosphorylation and activation (Sánchez-Sanz *et al*, 2016). Interestingly, mass spectrometry analysis of MST2 eluted fractions from HeLa lysates identified confidently the interaction between MST2 Lamin A/C (Figs 2A, EV2A) and suggested a potential interaction with Lamin B1. Lamin A/C and B1 both contribute to NE integrity, with Lamin B constituting more stable filaments and Lamin A/C more responsible for stable rigidity. We further examined the cellular localisation of nuclear MST1/2 kinases by IF in HeLa cells (with methanol fixation) and we observed that only MST2, and not MST1, exhibited perinuclear distribution similar to RASSF1A, as shown with co-staining with Lamin B1 (Fig 2B). To explore the role of MST2 localisation at the NE in relation to RASSF1A, we next reduced MST2 or RASSF1A expression using siRNAs and observed substantial reduction in RASSF1A staining at the INM in the absence of MST2 whereas, RASSF1A knockdown did not affect MST2 NE distribution (Fig 2C). Overall these results suggest that MST2 mediates RASSF1A localisation at the INM via lamina association.

**Figure 2:**
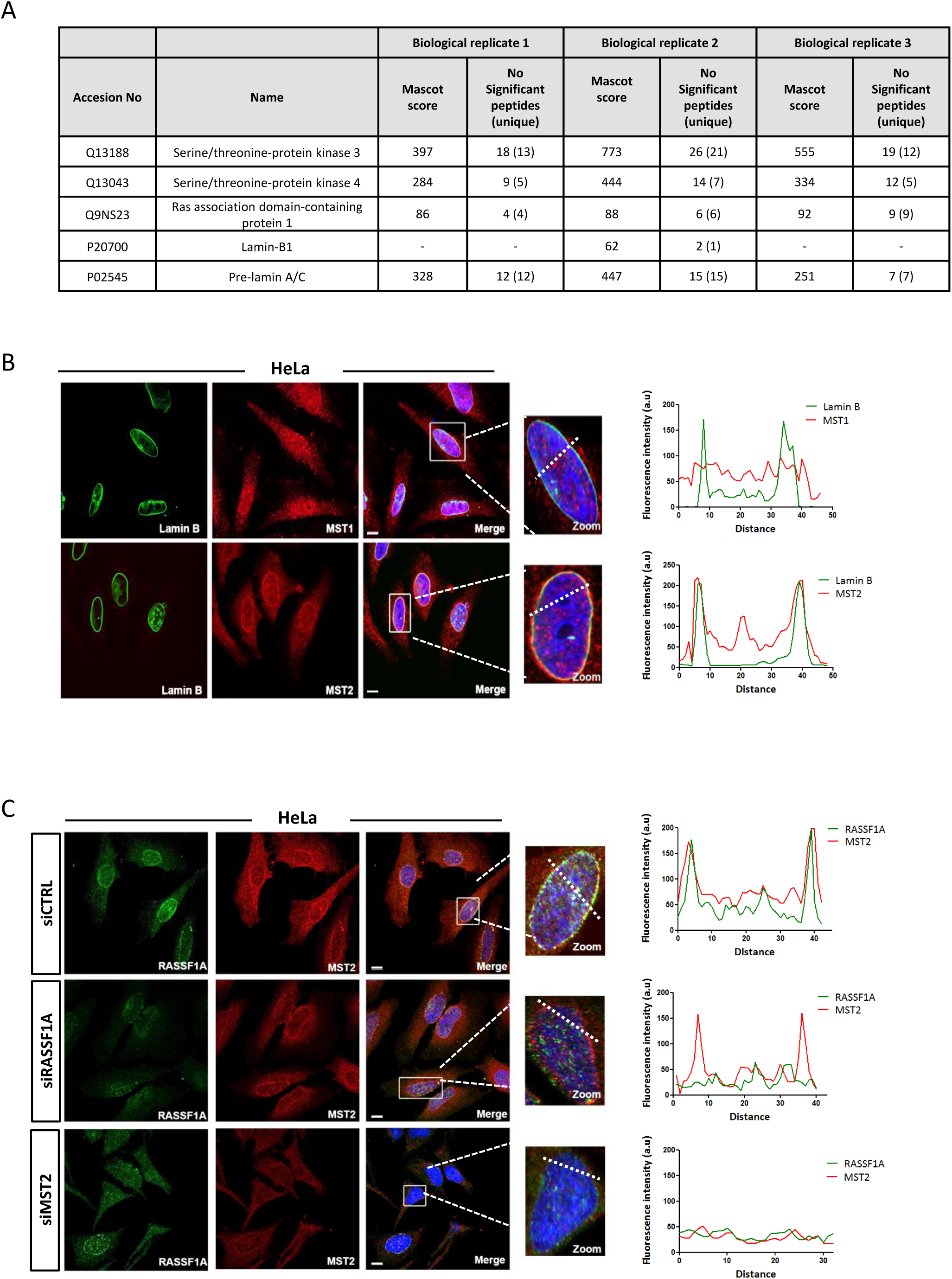
MST2 binds to nuclear pore complexes at the NE. A. List of proteins identified by LC-MS/MS in MST2 IP. The table includes the MASCOT score and number of significant peptides and unique peptides (in brackets) identified in each biological replicates after applying a 20 ion cut off and 1% FDR rate. Lamin B1 was identified only in 1 out of the 3 biological replicates with 1 unique peptide. However, interactions were verified by WB (Fig EV2A). B. Representative confocal images of MST1 (top) and MST2 (bottom) co-stained with the nuclear envelope marker Lamin B. Fluorescence intensity profile of Lamin B (green) and MST1 or MST2 (red) signals across the HeLa nuclei. Position of line scan indicated by the dashed white line. C. Immunofluorescence images of RASSF1A and MST2 in siCTRL, siRASSF1A and siMST2 treated cells. The graphs illustrate the fluorescence intensity profile of Lamin B (green) and MST2 (red) signals along the white lines shown in the merged panels.

### RASSF1A binds to XPO6 through its SARAH domain

We previously showed that RASSF1A interacts with RAN GTPase (Dallol *et al*, 2009), a key component in the regulation of nucleocytoplasmic transport. Given our data above showing the localisation of RASSF1A in the INM, we now sought to identify whether RASSF1A plays an active role in the process of RAN-dependent nuclear export. To test our hypothesis, we performed immunoprecipitation using extracts of HeLa cells and antibodies against exportin-1 (CRM1/XPO1), exportin-4 (XPO4), exportin-5 (XPO5), exportin-6 (XPO6) and exportin-7 (XPO7) (Fig 3A). Interestingly, we found that RASSF1A specifically interacts with XPO6, but not with CRM1/XPO1 which is the most conserved and responsible for export of a wide variety of cargoes (Fornerod *et al*, 1997).

**Figure 3:**
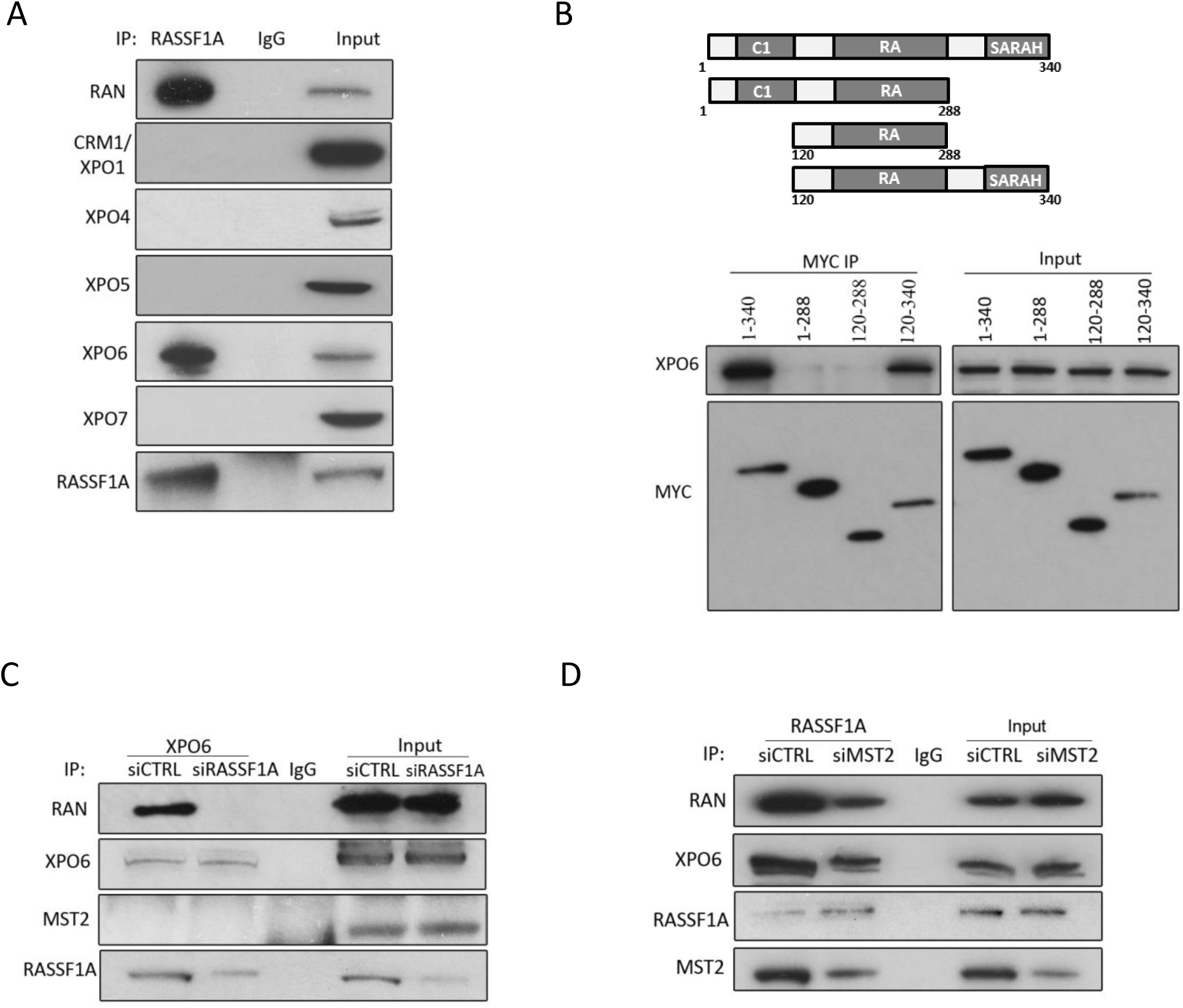
RASSF1A binds to XPO6 though its SARAH domain. A. Co-immunoprecipitation of endogenous RASSF1A with endogenous RAN, CRM1/XPO1, XPO4, XPO5, XPO6 and XPO7 from HeLa cell lysates, compared to the IgG control. B. *Upper:* Graphical representation of the domain structure of full-length RASSF1A and mutant constructs used for mapping RASSF1A/XPO6 interaction. Different domains are abbreviated as follows: SARAH, Salvador-RASSF-hippo domain; C1, N-terminal C1 type zinc fingers; RA, Ras binding domain. *Lower:* Western blot analysis of MYC immunoprecipitation from the indicated inputs from HeLa cells. Co-immunoprecipitation of endogenous XPO6 with RAN in siRASSF1A from HeLa cell lysates. C. Co-immunoprecipitation of endogenous XPO6 with endogenous RAN and RASSF1A in siRASSF1A HeLa cells. D. Co-immunoprecipitation of endogenous RASSF1A with endogenous XPO6 and RAN in siRNA-mediated knockdown of MST2 HeLa cells.

In order to determine the domain responsible for its interaction with XPO6, we exogenously expressed full-length MYC-RASSF1A and different RASSF1A-truncated mutants in HeLa cells. Whereas full-length MYC-RASSF1A (aa 1–340) and MYC-RASSF1A (aa 120–340) could specifically precipitate XPO6, no signal was detected with mutants lacking the SARAH domain, MYC-RASSF1A (aa 1–288) and MYC-RASSF1A (aa 120–288) (Fig 3B). Overall, these data demonstrate that the SARAH domain of RASSF1A mediates the interaction with XPO6. We further investigated the role of RASSF1A on the association of RAN to XPO6. Most strikingly, RASSF1A seemed to be required for the formation of XPO6/RAN complex, because siRNA-mediated knockdown of RASSF1A concomitantly decreased association between XPO6 and RAN (Fig 3C). Notably, XPO6/RAN interaction with RASSF1A does not include MST2, suggesting that this complex may be recruited to the NE via the RASSF1A-MST2 interaction. We next depleted MST2 expression using siRNA and observed decreased complex formation of RASSF1A with both XPO6 and RAN (Fig 3D) suggesting that failure to interact with MST2 and the NE disrupts XPO6/RAN stability.

### RASSF1A is involved in actin and profilin nuclear export process

XPO6 mediates specifically the export of actin and profilin complexes out of the nucleus (Stuven *et al*, 2003). We next determined whether RASSF1A plays a role in the XPO6-dependent nuclear export. Interestingly, subcellular nuclear/cytoplasmic fractionation of HeLa cell lysates clearly demonstrated elevated levels of profilin and actin in the nucleus upon siRNA-mediated silencing of RASSF1A, compared to cells treated with control siRNA, suggesting an impaired nuclear export (Fig 4A). Nuclear retention of profilin in HeLa cells treated with siRNA against RASSF1A was further demonstrated by immunofluorescence (Fig 4B). Interestingly, *RASSF1A* gene silencing does not affect the expression levels of XPO6 (Fig EV3A). Additionally, to rule out the possibility that nuclear actin accumulation resulted from impaired nuclear import receptor levels, we next examined the protein expression of importin-9 (IPO9), which is required for actin translocation to the nucleus (Dopie *et al*, 2012), in the absence of RASSF1A. Western blotting showed no change in the expression of IPO9 upon depletion of RASSF1A, compared to the siCTRL-transfected cells (Fig EV3A). The significance of RASSF1A for XPO6-mediated export process is also evident in MDA-MB-231 cells that lack RASSF1A expression. Strikingly, forcing the expression of *RASSF1A* either by transiently transfecting MDA-MB-231 cells with a plasmid encoding *RASSF1A* or by treating cells with the demethylating agent 5’-aza-dC, led to a decrease in actin and profilin nuclear retention (Fig 4C, 4D). Taken together, these data strongly indicate that RASSF1A governs the export of actin and profilin out of the nucleus.

**Figure 4:**
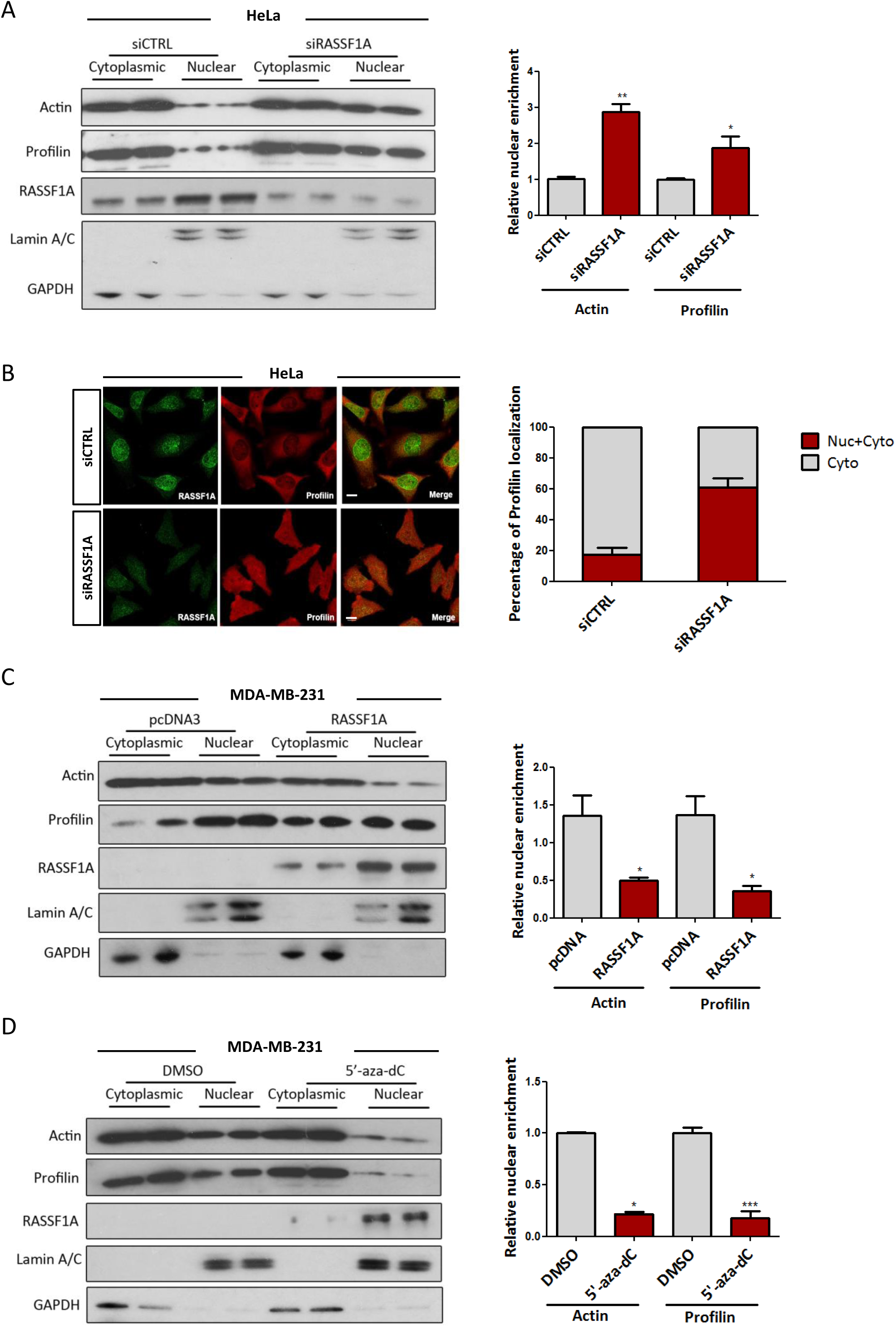
RASSF1A is involved in actin and profilin nuclear export process. A. HeLa cells treated with control or *RASSF1A* siRNA were fractionated into cytoplasmic and nuclear extracts. Lysates from each fraction were probed for expression levels of actin and profilin alongside GAPDH (as a marker of the cytoplasmic fraction) and Lamin A/C (as a marker of the nuclear fraction). *Right:* Quantification of nuclear actin and profilin was done relative to the control Lamin A/C. *P <0.05, **P<0.01 B. Immunofluorescence images of profilin in control and RASSF1A siRNA transfected HeLa cells. *Right:* The Profilin localisation was scored as nuclear/cytoplasmic or predominantly cytoplasmic in approximately 100 cells. C. Western blot analysis of actin, profilin, GAPDH and Lamin A/C levels in nuclear and cytoplasmic fractions of MDA-MB-231 cells treated with control pcDNA3 or *RASSF1A* vector. The graph shows the nuclear levels of actin and profilin in MDA-MB-231 cells expressing RASSF1A. *P <0.05 D. Western blot analysis of actin, profilin, GAPDH and Lamin A/C levels in nuclear and cytoplasmic fractions of MDA-MB-231 cells treated with DMSO or 5’ Aza. The graph shows the nuclear levels of actin and profilin in MDA-MB-231 cells expressing RASSF1A. *P <0.05, ***P<0.001

### Loss of RASSF1A expression alters MRTF-A/SRF axis

Previous studies showed that nuclear actin has been linked to gene transcription (Visa & Percipalle, 2010; Kapoor & Shen, 2014; Grosse & Vartiainen, 2013), downregulating the expression of *MYL9, ITGB1* and *PAK1* (Sharili *et al*, 2016). Therefore, we measured mRNA levels of these nuclear actin-regulated genes and show that these genes were significantly reduced following siRNA-mediated knockdown of RASSF1A in comparison to control (siCTRL) HeLa cells (Fig 5A). Furthermore, mRNA encoding the transcription factor *OCT4*, which is known to be activated by nuclear actin (Yamazaki *et al*, 2015), was elevated in HeLa cells depleted of RASSF1A (Fig 5A). It is well established that depletion of IPO9 promotes the export of actin from the nucleus (Dopie *et al*, 2012). As expected, the expression of actin in nuclear extracts from HeLa cells treated with siRNA against IPO9 was decreased and prevented accumulation even in the absence of export via RASSF1A silencing (Fig EV3B). Moreover, the increased levels of *MYL9, ITGB1, PAK1* and *OCT4* mRNA upon RASSF1A silencing were rescued in HeLa cells co-depleted of IPO9 (Fig 5A).

**Figure 5:**
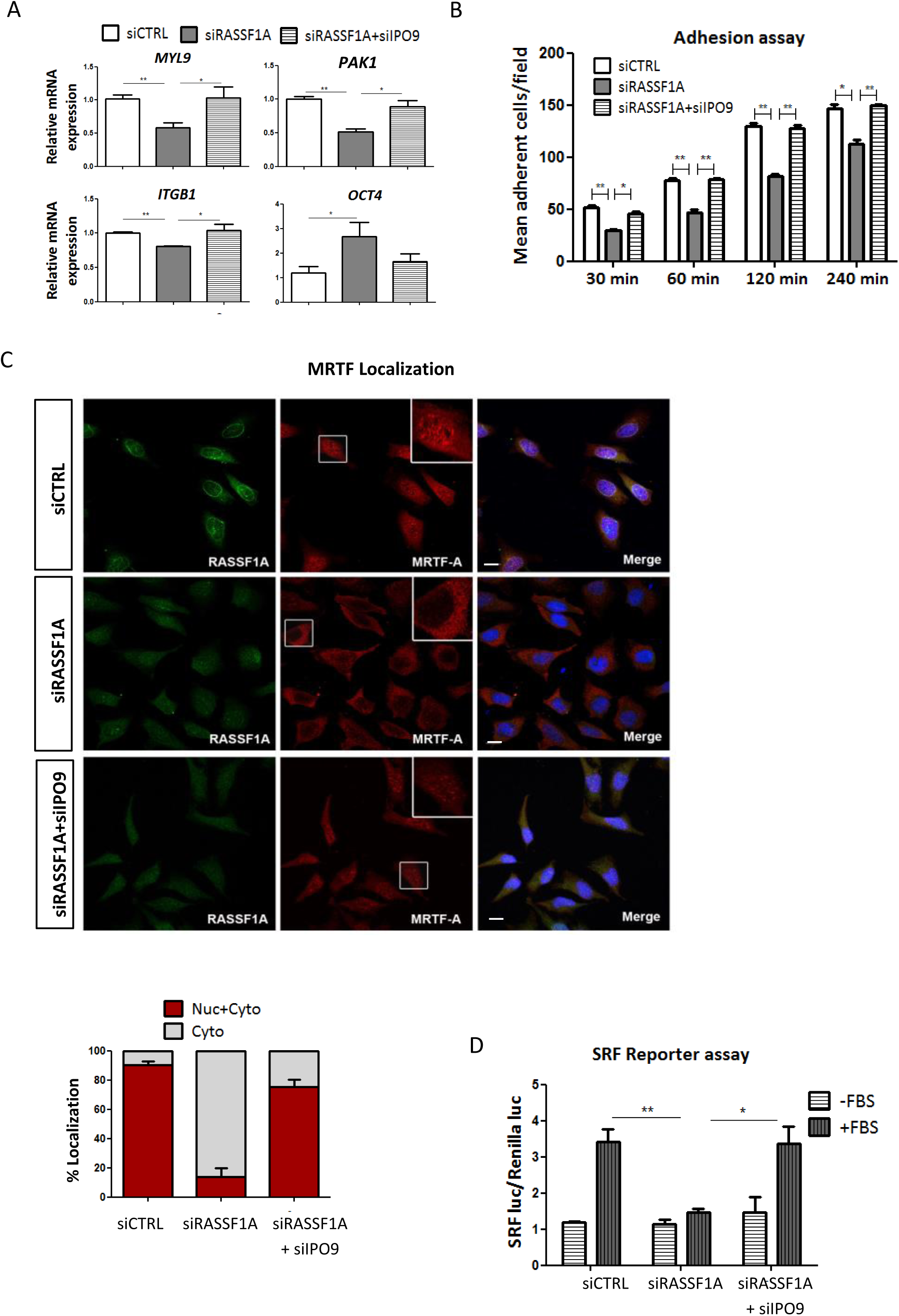
Loss of RASSF1A expression alters MRTF-A/SRF axis. A. QRT-PCR validation of selected genes known to be affected by the levels of nuclear actin. Expression levels of *MYL9, ITGB1, PAK1* and *OCT4* from HeLa cells treated either with siRASSF1A or siRASSF1A+siIPO9 are relative to *GAPDH* and normalised to siCTRL cells. Data represent the mean ± SEM of 3 replicates. *P <0.05, **P<0.01 B. Adhesion assay. siRNA against *RASSF1A* significantly decreased HeLa cells adhesive rate at all the determined time points compared to control. The cells were cultured for 48 h before harvesting and reseeding for 1 h on 96 well-plates coated with FN. Data represent the mean±SEM of 3 replicates. **P=0.003, ***P<0.0001 C. Representative immunofluorescence images of MRTF-A localisation in siRASSF1A and siRASSF1A+siIPO9. *Lower:* The MRTF-A localisation was scored as nuclear/cytoplasmic or predominantly cytoplasmic in 100-200 cells. Scale bars are 10 μm. D. Luciferase assay of SRF dependent promoter in cells transfected with siCTRL, siRASSF1A orsiRASSF1A+siIPO9 following stimulation with 10% FBS for 5 h. Data are expressed as SRF luciferase activity relative Renilla control and represent the mean ± SEM of 3 replicates. *P < 0.05 for siRASSF1A+siIPO9 vs siRASSF1A (+ FBS), **P < 0.01 for siRASSF1A vs siCTRL (+FBS)

We next tested whether increased nuclear actin levels arising from RASSF1A depletion have functional consequences for the cells. Since multiple nuclear actin-regulated genes encode for known regulators of cell adhesion (Sharili *et al*, 2016), we then assessed whether these cellular functions are affected in RASSF1A-depleted cells. In accordance with our qRT-PCR data, we showed that HeLa cells lacking RASSF1A exhibited a significantly decreased number of adhesive cells (Fig 5B). Accordingly, restoring actin levels in the nucleus by silencing IPO9 in the absence of RASSF1A was sufficient to rescue this effect (Fig 5B). Collectively, our data show that RASSF1A has an indirect effect on the expression of genes involved in cell adhesion processes, via regulation of actin levels within the nucleus.

Nuclear actin plays a key role in the regulation of the localisation and activity of Myocardin-related transcription factor A (MRTF-A), a co-activator of the transcription factor serum response factor (SRF), which regulates the expression of many cytoskeletal genes (Sotiropoulos *et al.*, 1999; Miralles *et al.*, 2003; Vartiainen *et al.*, 2007; Ho *et al.*, 2013). We next hypothesized that MRTF-A localisation and transcriptional activity of SRF could both be affected by RASSF1A protein levels. Notably, we observed that MRTF-A resides mostly in the cytoplasm in the absence of RASSF1A whereas it is located in both nucleus and cytoplasm in control cells (Fig 5C). Increased nuclear actin levels exclude MRTF-A from the nucleus and block SRF-dependent gene transcription (Vartiainen *et al*, 2007a; Posern *et al*, 2002). Accordingly, we showed decreased mRNA levels of the *SRF* gene together with decreased FBS-stimulated SRF-reporter activity in cells transfected with siRNA against RASSF1A (Figs 5D, EV3C). Silencing IPO9 restored MRTF-A nuclear localisation and SRF expression in RASSF1A-depleted cells (Figs 5C, 5D). These results demonstrate a correlation between RASSF1A expression and SRF axis regulation, via actin localisation.

We next analysed *SRF* gene expression from clinical data available from The Cancer Genome Atlas (TCGA) database with the cBioportal tool (http://www.cbioportal.org). *RASSF1A* methylation is widely appreciated to correlate with adverse prognosis and we find a significant correlation between this epigenetic event and *SRF* mRNA levels indicating, that loss of nuclear actin regulation and MRTF/SRF is likely to contribute to clinical parameters (Fig 6A, top). In line with a role in tumour suppression, *SRF* levels are significantly reduced in invasive breast cancer compared to control normal tissues (*Wilcoxon p=1.164e*^*-20*^; Fig 6B, bottom). Additionally, Breast Invasive Carcinomas (TCGA) display higher *SRF* mRNA expression in tumours with high levels of *RASSF1A* transcript (*p<0.0001*), which also shows significant linear correlation across the dataset (*R=0.28*) (Fig 6B). We find a matching association of *RASSF1A* and *SRF* in Hepato-Cellular Carcinoma (TCGA) (*p<0.0001*), again with significant linear correlation of *RASSF1/SRF* transcripts (*R=0.29*) (Fig 6C) and further correlations with bladder and colorectal cancer (Fig EV4). Notably, *SRF* mRNA levels were enriched in the RASSF1^high^ individuals in four distinct cancer types (Figs 6 and EV4). Finally, we examined the Breast and Liver cancer survival statistics in the TCGA and we found that patients with low SRF expression have a poor prognosis for overall survival (Breast HR = 0.8 (0.72 – 0.89) Logrank *P =5e*^*-5*^; HCC HR = 0.67 (0.47 – 0.96) Logrank *P =0.027*). Overall our data show a correlation of *RASSF1* and *SRF* mRNA expression and SRF levels associate with poor survival.

**Figure 6:**
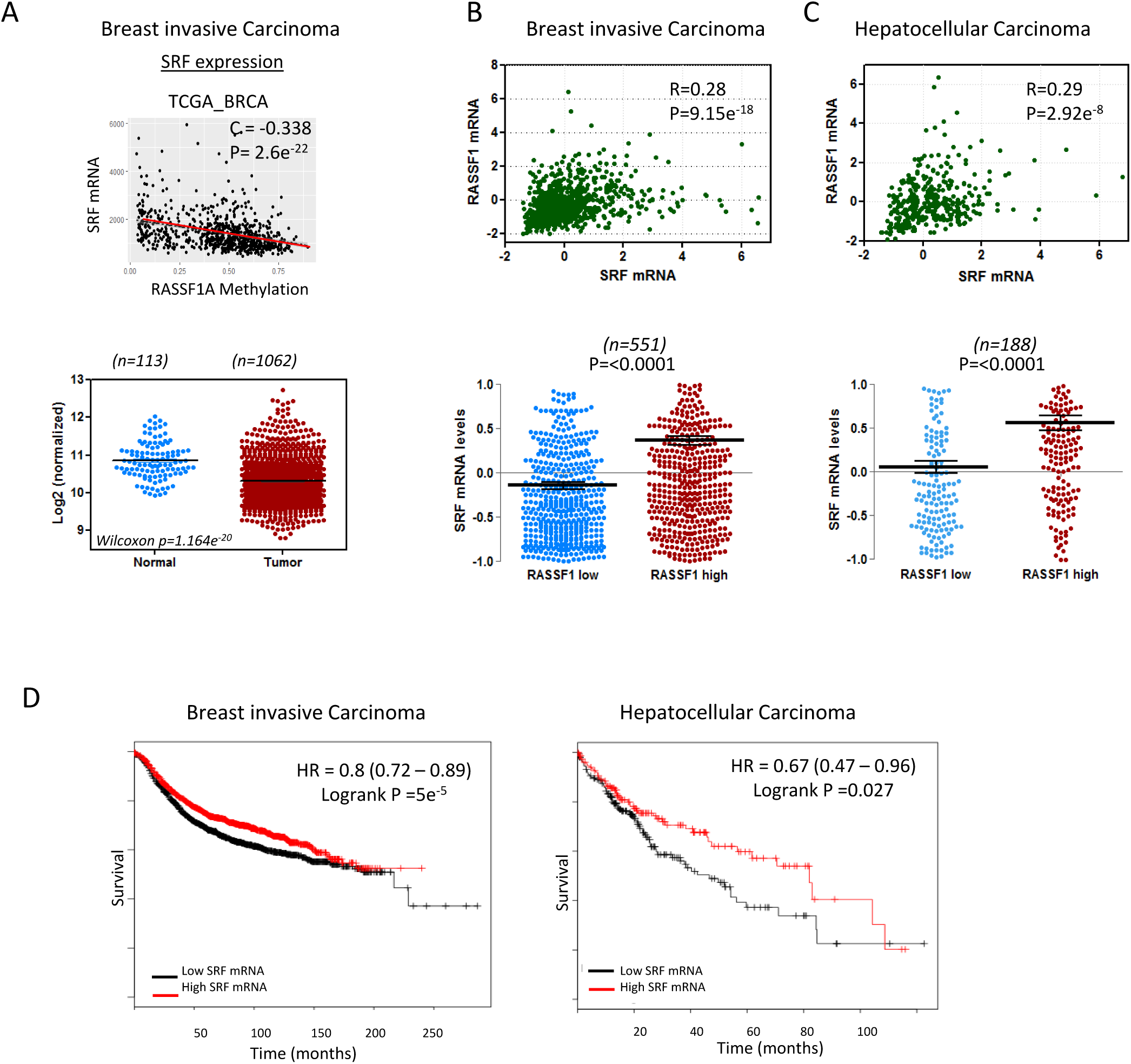
Correlation of RASSF1 and SRF expression. A. Upper: Scatter plot of RASSF1 methylation levels to SRF mRNA expression in breast invasive carcinoma (TCGA, Provisional, n=960). Lower: SRF expression (TCGA, Illumina HiSeq RNAseq) in normal and BRCA patients. B. Upper: Scatter plot of RASSF1 mRNA to SRF mRNA levels in breast invasive carcinoma (TCGA, Provisional, n=960). Values are given in (RNA Seq V2 RSEM). Lower: Differences in SRF mRNA expression of samples that express low and high levels of RASSF1 mRNA. C. Upper: Scatter plot of RASSF1 mRNA to SRF mRNA levels in hepatocellular carcinoma (TCGA, Provisional, n=360 samples). Values are given in (RNA Seq V2 RSEM). Lower: Differences in SRF mRNA expression of samples that express low and high levels of RASSF1 mRNA. D. Survival analysis of BRCA (n=3951) and HCC (n=364) patient samples estimated by Kaplan Meier survival curve with high (red) and low (black) expression of SRF.

## Discussion

Actin constantly shuttles between the cytoplasm and the nucleus using an active transport mechanism and proper balance of nuclear and cytoplasmic actin pools is necessary to maintain cellular homeostasis (Dingová *et al*, 2009). Nuclear actin exported from the nucleus by XPO6, independently of the general export receptor CRM1, and it is imported by IPO9 (Dopie *et al*, 2012; Stuven *et al*, 2003). As for most export processes, XPO6 also requires RAN GTPase to export actin monomers in complex with profilin from the nucleus (Stuven *et al*, 2003). We found that lack of RASSF1A, a phenomenon commonly observed in human tumours, leads to accumulation of actin within the nucleus due to defective nuclear export. The direct interaction of RASSF1A with XPO6, a member of the importin-β superfamily of transport receptors, is required for the association with RAN. The XPO6 complex does not appear to include MST2 suggesting that the recruitment of RASSF1A to MST2 at the NE potentially displaces XPO6 binding to the SARAH domain (Dittfeld *et al*, 2012) and this is important for the active participation in actin-profilin nucleo-cytoplasmic shuttling. Reduction in cytoplasmic actin was not evident as a consequence of the defective export, most likely because of the higher actin concentration in the cytoplasm (Fig 4A). Nuclear actin is a key regulator of transcription and it is required for all three RNA polymerases (Pol I, II and III) (Qi *et al*, 2011; Philimonenko *et al*, 2004). Previous reports showed that expression of nuclear actin negatively regulates multiple genes and results in altered expression of approximately 2000 genes (Yamazaki *et al*, 2015; Sharili *et al*, 2016). Loss of *RASSF1A* expression alters gene expression and impacts cellular processes, such as cell adhesion. Thus, RASSF1A appears to contribute to the transcriptional activity of the cell and further to its phenotypic changes through its ability to modulate nuclear actin levels. Although nuclear actin levels have not been extensively studied in cancer cells, abnormal subcellular localisation of certain ABPs (actin-binding proteins) is associated with the development carcinogenesis processes (Honda, 2015; Khurana *et al*, 2011; Savoy & Ghosh, 2013; Loy *et al*, 2003; Velkova *et al*, 2010; Patnaik *et al*, 2016). A recent study showed that the nuclear actin levels regulated by XPO6 are pivotal for cell proliferation and quiescence and that preventing actin export from the nucleus contributes to malignant progression (Fiore *et al*, 2017). Of note, low XPO6 expression and therefore nuclear actin accumulation correlates with poor survival of breast cancer patients (Fiore *et al*, 2017).

Mechanotransduction signaling is essential for a broad range of biological functions such as embryogenesis, cell migration, metastasis and epithelial-mesenchymal transition. MRTF-A is a mechanosensitive transcription co-factor which, together with SRF, controls a variety of genes involved in actin cytoskeletal remodelling and growth factor response. MRTF–SRF activation is responsible for immediate transcriptional response to mechanical stimuli (Cui *et al*, 2015) and perturbed MRTF-SRF signaling axis might partially explain the pathogenesis of certain human diseases (Ho *et al*, 2013). We found that the SRF-driven gene transcription activity in response to serum stimulation was severely abrogated in RASSF1A depleted cells (Fig 4D). In addition, we also found that RASSF1 expression correlates with *SRF* expression in a variety of cancers (Fig 6). Although there is currently lack of evidence, altered MRTF-SRF signaling may explain the aberrant mechanosensitivity observed often in cancer cells (Tang *et al*, 2012; Ciasca *et al*, 2016; Ghosh *et al*, 2008).

Epigenetic silencing of the *RASSF1A* gene has been frequently associated with poor patient outcome across all solid malignancies (Grawenda & O’Neill, 2015). Several studies have identified functional roles for RASSF1A mediated tumour suppression that can explain these clinical associations, e.g., hippo pathway regulation (Hamilton *et al*, 2009; Matallanas *et al*, 2007), apoptosis (Vos *et al*, 2000; Baksh *et al*, 2005), differentiation (Papaspyropoulos *et al*, 2018), the cytoskeleton (Liu *et al*, 2003; Dallol *et al*, 2004; Vlahov *et al*, 2015) and DNA damage/repair (Pefani *et al*, 2014, 2018). However, the widespread prognostic associations imply a perturbation of a biological process that simultaneously contributes to these diverse mechanisms but this remains unexplained. Intriguingly, nuclear actin levels have also been described to contribute to the regulation of hippo pathway (e.g. SRF regulation of YAP) (Foster *et al*, 2017; Sen *et al*, 2015), apoptosis (Sharili *et al*, 2016), differentiation (Sen *et al*, 2015, 2017; Xu *et al*, 2010) and DNA damage/repair (Belin *et al*, 2015; Yuan & Shen, 2001). Altogether, our findings indicate that loss of RASSF1A expression results in failure to export nuclear actin, suggesting that both regulatory processes are linked and that the clinical data associated with *RASSF1* methylation involves deregulated MRTF/SRF.

## Methods

### Tissue culture and cell treatments

HeLa, MEFs, HEK293, RPE-1 and MDA-MB-231 cells were cultured in complete DMEM supplemented with 10% foetal bovine serum in 5% CO2 and 20% O2 at 37°C. HeLa cells were purchased from Cancer Research UK, London, or LGC Promochem (ATCC). Inhibitors for ATR kinase activity (VE-821), ataxia telangiectasia mutated (ATM) kinase activity (KU-55933) were used at a concentration of 10 μM. 10 µM 5-aza-2′-deoxycytidine (5’-aza-dC; Sigma) was added to the medium, and the cells were incubated for 72 h. Due to its chemical instability, 5’-aza-dC was added to the fresh medium every 24 h.

### Real-time PCR primers

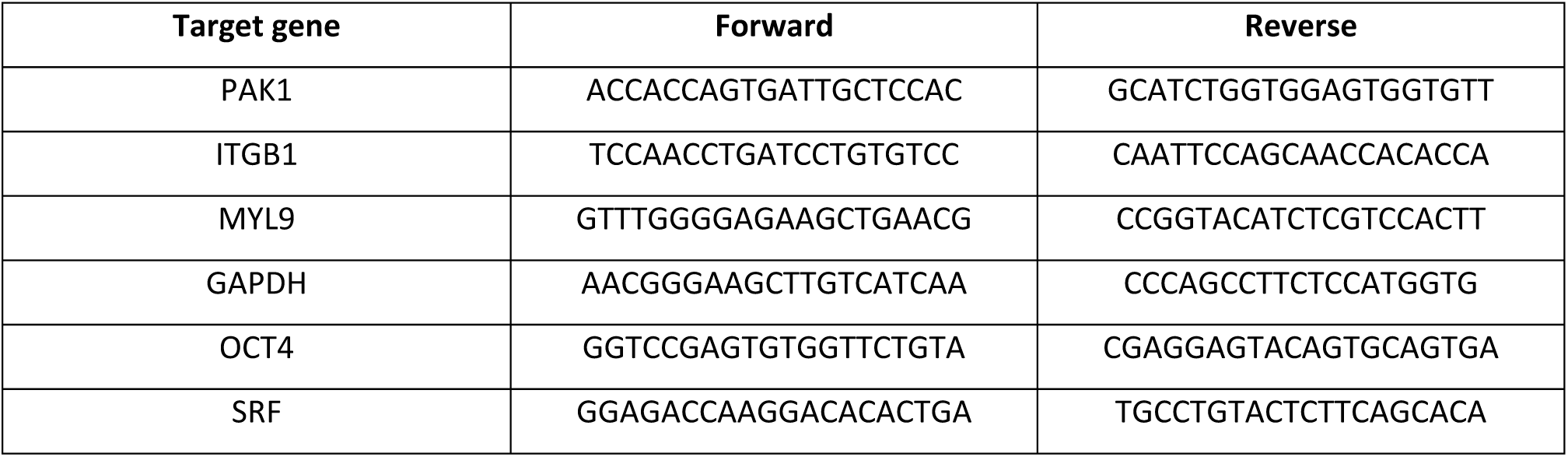

### siRNA oligonucleotides

RNA interference was carried out with 100 nM siRNA for 48 h using Lipofectamine 2000 (Invitrogen) according to manufacturer’s instructions. The oligos used were: siMST2: siGENOME smartpool: M-004874-02 (Dharmacon), siRASSF1A: GACCUCUGUGGCGACUU, siLaminA/C: sc35776 (Santa Cruz Biotechnology, Inc), siNUP133: sc60035 (Santa Cruz Biotechnology, Inc). siRNA against Lusiferase with the sequence GCCAUUCUAUCCUCUAGAGGAUG was used as control.

### Plasmids

RASSF1A truncation mutants, MYC-RASSF1A 1-288, MYC-RASSF1A 120-340, MYC-RASSF1A120-288 were created by PCR using MYC-tagged RASSF1A as template, as previously shown (Pefani *et al*, 2016). MDA-MB-231 cells were transiently transfected with 1 µg of RASSF1A DNA or the empty pcDNA 3.0 plasmid using Lipofectamine 2000 (Invitrogen).

### Antibodies

The following antibodies were used in this study: RASSF1A (Atlas, HPA040735), MST1 (Sigma, WH0006789M1), MST2 (Abcam, ab71960), XPO6 (Bethyl, A301-205A), CRM1 (Cell signaling, 46249), XPO4 (Novus Biologicals, NB100-56495), XPO5 (sc-166789), XPO7 (sc-390025), Lamin A/C (Cell signaling, 4777), Lamin B (Abcam, ab16048), Profilin (Novus Biologicals, NBP2-02577), IPO9 (Abcam, ab52605), Myc-Tag (JBW301, Millipore, 05-724), GAPDH (Cell signaling, 97166), Actin (sc-1616, Santa Cruz), RAN (Santa Cruz Biotechnology, sc58467), BrdU (abcam, ab6326), p EV1 31-RASSF1A custom-made (Hamilton *et al*, 2009).

### Quantitative real-time PCR analysis

RNA extraction, reverse transcription and qPCR were performed using the Ambion PowerSYBR Green Cells-to-CT kit following manufacturer’s instructions in a 7500 FAST Real-Time PCR thermocycler with v2.0.5 software (Applied Biosystems). The mRNA expression level of each gene relative to GAPDH was calculated using the ΔΔCt method. All experiments were performed in triplicates.

### Protein extraction and immunoblotting

Cytosolic and nuclear fractions were prepared using the NE-PER Nuclear and Cytosolic Extraction Reagents (Thermo Fisher Scientific) according to the manufacturer’s instructions. Sample protein content was determined by the BiCinchoninic Acid Assay protein assay (Thermo Fisher Scientific). Extracts were analyzed by SDS-PAGE using a 10% Bis–Tris Nu-PAGE gels (Invitrogen) and transferred onto PVDF membranes (Millipore). Subsequent to being washed with PBS containing 1% Tween 20 (PBS-T), the membranes were blocked in 5% bovine serum albumin (BSA) in TBS-T for 1 h at RT, then incubated with the primary antibody overnight at 4°C. The membranes were incubated with HRP-conjugated secondary antibodies for 1h at room temperature and exposured to X-ray film (Kodak). ImageJ software (NIH) was used for the quantification of the bands. All bands were normalised against the loading controls.

### Immunofluorescence

HeLa cells were grown on coverslips, washed with PBS and fixed with MeOH at 20°C for 20 min. Where indicated, cells were instead fixed in 4% paraformaldehyde and permeabilised either with 0.5% Triton X-100 at room temperature for 5 min or with 40 μg/ml digitonin on ice for 2 min. Coverslips were then incubated with primary antibody for 2 h in PBS with 3% BSA at room temperature. For the 5-Fluorouridine incorporation, cells were incubated with 2 mM 5-FUrd (Sigma) for 20 min, fixed and stained with rat anti-BrdU. Cells were washed with PBS and incubated with secondary anti-rabbit and/or anti-mouse or anti-rat IgG conjugated with Alexa-Flour 488 or Alexa-Flour 568 (Molecular Probes) for 1 h at room temperature. Samples were washed and mounted on coverslips with mounting medium containing DAPI (Invitrogen). Cells were analysed using LSM780 (Carl Zeiss Microscopy) confocal microscope. Fluorescence intensity plots were generated using ImageJ software (NIH).

### Duolink PLA

Duolink PLA was performed using the Duolink Red Kit (Sigma). In summary, cells were seeded at a density of 4 × 10^5^ cells per well of a 12-well plate. Cells were fixed in 4% formaldehyde after 24 hours, permeabilised and blocked following the manufacturer’s instructions. Following incubation with primary antibodies, coverslips were washed and incubated with a mixture of the PLUS/MINUS Duolink^®^ PLA probes (OlinkBioscience) for 1 hour at 37 °C. Coverslips were then washed and detection of signal was conducted using the Duolink^®^ detection reagent kit (red) (OlinkBioscience). Coverslips were first incubated with a ligation-ligase mixture for 30 min at 37 °C, washed and further incubated with an amplification-polymerase mixture for 100 min at 37 °C. Subsequently, coverslips were washed and imaged using the Zeiss microscope system.

### Immunoprecipitation

Cells were lysed in lysis buffer (50 mM Tris-HCl pH 7.5, 0.15 M NaCl, 1mM EDTA, 1% NP-40, 1 mM Na_3_VO_4_, complete proteinase inhibitor cocktail (Roche)), and incubated with 20 μl protein G Dynabeads (Invitrogen) and 2 μg of the indicated antibodies at for 2 h at 4°C. Pelleted beads were collected in sample buffer NuPAGE LDS (Thermo Fisher Scientific) with 200 mM DTT and subjected to SDS-PAGE and immunoblotting.

### Mass spectrometry

HeLa cells were lysed in 1% NP-40 lysis buffer (150 mM NaCl, 20 mM HEPES, 0.5 mM EDTA) containing complete protease and phosphatase inhibitor cocktail (Roche). 10 mg of total protein lysate were incubated for 3 h with protein A Dynabeads (Invitrogen) and 10 μg of MST2 antibody (ab52641) or rIgG at 4°C. MST2 immunoprecipitates were eluted off the beads in a low pH glycine buffer. The eluted fractions were sequentially incubated with DTT (5mM final concentration) and Iodacetamide (20mM final concentration) for 30min at room temperature in the dark, before proteins were precipitated with methanol/chloroform. Protein precipitates were reconstituted and denatured with 8M urea in 20mM HEPES (pH 8). Samples were then further diluted to a final urea concentration of 2M using 20Mm HEPES (pH8.0) before adding immobilised trypsin for 16h at 37°C (Pierce 20230); (Montoya et al. 2011). Trypsin digestion was stopped by adding 1% TFA (final concentration) and trypsin beads removed by centrifugation. Tryptic peptides solution was desalted by solid phase extraction using C18 Spintips (Glygen LTD) and dried down.

Dried tryptic peptides were re-constituted in 15 µl of LC-MS grade water containing 2% acetonitrile and 0.1% TFA. Seven percent of the sample was analysed by liquid chromatography-tandem mass spectrometry (LC_MS/MS) using a Dionex Ultimate 3000 UPLC coupled to a Q-Exactive mass spectrometer (Thermo Fisher Scientific). Peptides were loaded onto a trap column (PepMapC18; 300µm × 5mm, 5µm particle size, Thermo Fischer) for 1 min at a flow rate of 20 μl/min before being chromatographic separated on a 50cm-long EasySpray column (ES803, PepMAP C18, 75 µm × 500 mm, 2 µm particle, Thermo Fischer) with a gradient of 2 to 35% acetonitrile in 0.1% formic acid and 5% DMSO with a 250 nL/min flow rate for 60 min (Chung et al. 2016). The Q-Exactive was operated in a data-dependent acquisition (DDA) mode to automatically switch between full MS-scan and MS/MS acquisition. Survey-full MS scans were acquired in the Orbitrap mass analyser over an m/z window of 380 – 1800 and at a resolution of 70k (AGC target at 3e6 ions). Prior to MSMS acquisition, the top 15 most intense precursor ions (charge state >=2) were sequentially isolated in the Quad (m/z 1.6 window) and fragmented in the HCD collision cell (normalised collision energy of 28%). MS/MS data was obtained in the Orbitrap at a resolution of 17500 with a maximum acquisition time of 128ms, an AGC target of 1e5 and a dynamic exclusion of 27 seconds.

The raw data were searched against the Human UniProt-SwissProt database (November 2015; containing 20268 human sequences) using Mascot data search engine (v2.3). The search was carried out by enabling the Decoy function, whilst selecting trypsin as enzyme (allowing 1 missed cleavage), peptide charge of +2, +3, +4 ions, peptide tolerance of 10 ppm and MS/MS of 0.05 Da; #13C at 1; Carboamidomethyl (C) as fixed modification; and Oxidation (M) and Deamidation (NQ) as a variable modification. MASCOT outputs were filtered using an ion score cut off of 20 and a false discovery rate (FDR) of 1%. The mass spectrometry raw data included in this paper had been deposited to the ProteomceXchange Consortium via the PRIDE partner repository with the dataset identifier PRIDE: PXD11517.

### Cell adhesion assay

HeLa cells were transfected with siRNA against RASSF1A or RASSF1A and IPO9 or control and were collected after 48 h. (1×10^5^/ml, 0.5 ml) were re-plated on fibronectin-coated 24-well plate in triplicate. After attachment for the predetermined time (30 min, 60 min and 90 min), the plate was washed to remove the non-adherent cells. The number of adherent cells from 4 random fields of the well was counted microscopically.

### SRF reporter assay

Cells were transfected with the SRF luciferase reporter (pGL4.34) using Lipofectamine 2000. pRL Renilla luciferace vector was used as an internal control. Following DNA transfection, cells were rinsed and cultured for 24 hours before treating with FBS. Luciferase assays were carried out in 24-well plates (n=3 wells); cells were treated for 7hours with FBS, then harvested and analyzed using the dual luciferase assay (Promega).

### Correlation analysis

The TCGA data were downloaded from the cBioPortal for Cancer Genomics (http://www.cbioportal.org). The association of RNA expression of *RASSF1* and *SRF* was studied in breast invasive carcinoma (TCGA, Provisional), hepatocellular carcinoma (TCGA, Provisional), bladder cancer (TCGA, Cell 2017) and colorectal adenocarcinoma (TCGA, Nature 2012). Methylation 450K data and gene expression data were downloaded from TCGA (BRCA). RASSF1A methylation levels for each patient were determined as the average beta value across all probes within the RASSF1A CpG island (<u>chr3:50377804-50378540</u>). Pearson correlation and Scatterplots were drawn using R (Ver 3.4). Survival analyses from 3951 BRCA samples and 364 liver HCC patient samples were performed using the Kaplan-Meier Plotter tool with the patients being splitted by median (Nagy *et al*, 2018).

### Statistics

For all experiments, statistical analysis was carried out using a Student’s t test. All data are expressed as mean ± SEM.

## Author contributions

M.C. conceived the project and designed the research, performed experiments and analyzed data. D.E.P., D.P and M.E assisted with experiments. I.V and R.F performed the mass spectrometry analysis. E.O.N. and M.C. wrote the manuscript.

## Acknowledgements

This work was funded by Cancer Research UK A19277. Mass spectrometry analysis was performed at the Discovery Proteomics Facility (headed by Roman Fischer) which is part of the TDI MS Laboratory (led by Benedikt Kessler).

---

**Supplementary Figure 1.**
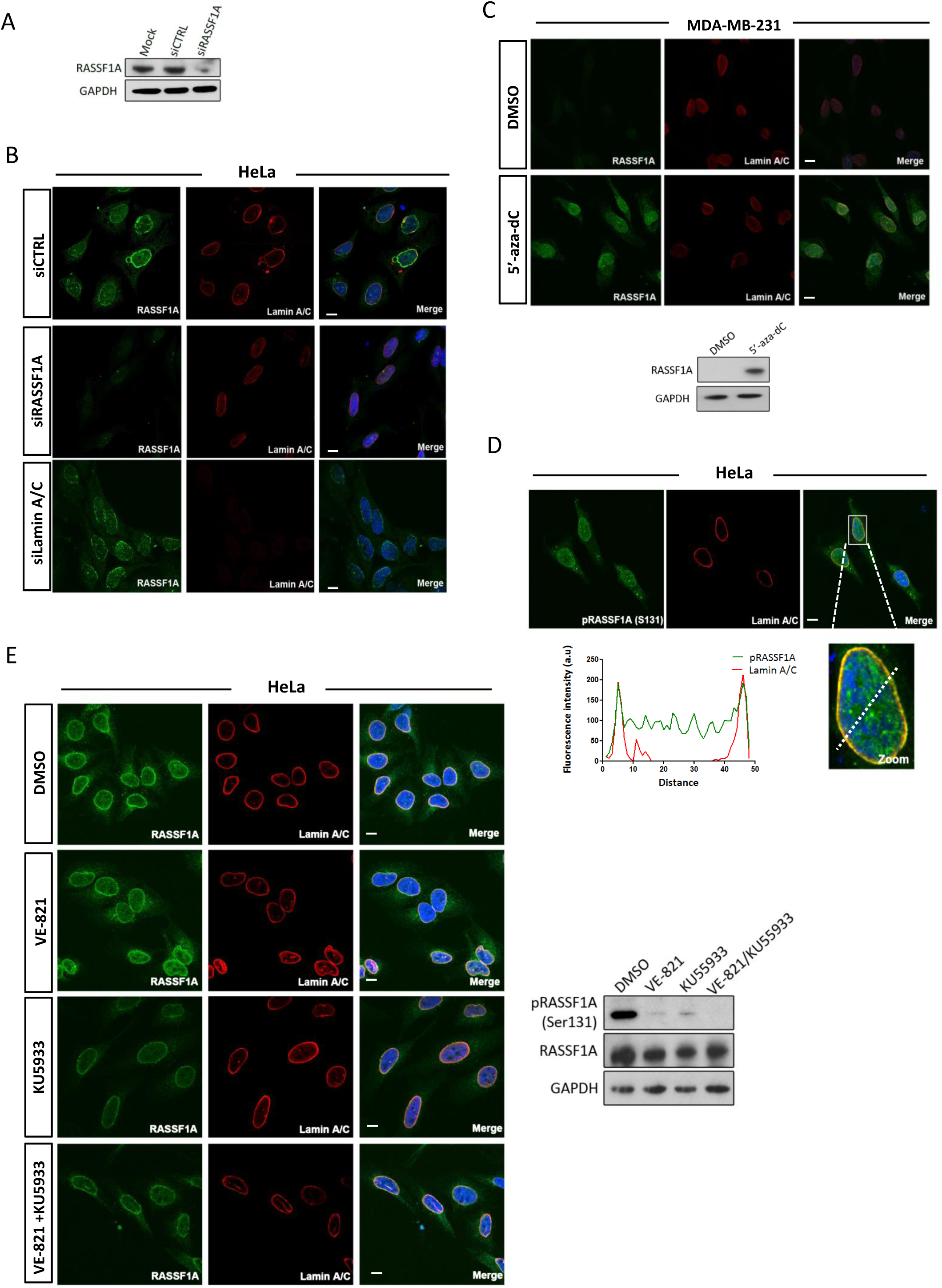
A. Assessment of RASSF1A protein levels in HeLa cells transfected with siRASSF1A. B. Representative confocal images of RASSF1A and Lamin A/C in siRASSF1A and siLamin A/C transfected HeLa cells. C. Immunofluorescence images of RASSF1A in MDA-MB-231 cells after treatment with DMSO or 5’-aza-dC demethylating reagent. Lower blot shows the expression levels of RASSF1A levels following treatment. D. Immunofluorescence detection of phosphorylated of phosphorylated RASSF1A (EV1 31). Fluorescence intensity profile of Lamin A/C (red) and pRASSF1A (EV1 31) (green) signals across the HeLa nuclei. Position of line scan indicated by the dashed white line. E. Immunofluorescence images of RASSF1A in HeLa cells treated with DMSO, ATR inhibitor VE821, ATM inhibitor KU5933 and the combination of both. Western blot (right) of RASSF1A protein levels following the corresponding treatments.

**Supplementary Figure 2.**
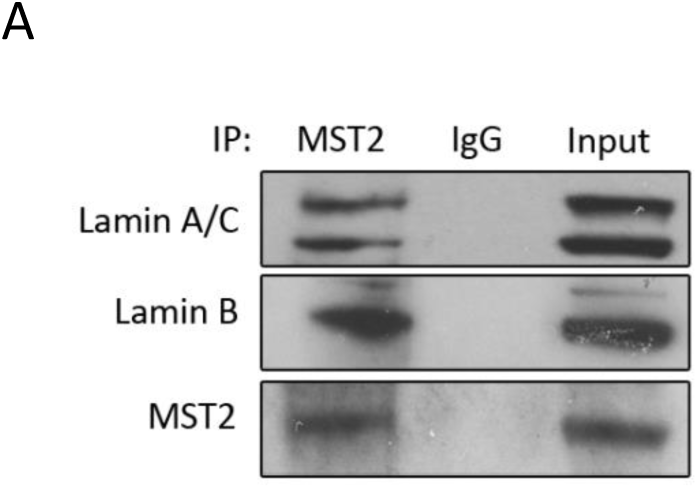
A. Co-immunoprecipitation of endogenous MST2 with Lamin B and Lamin A/C.

**Supplementary Figure 3.**
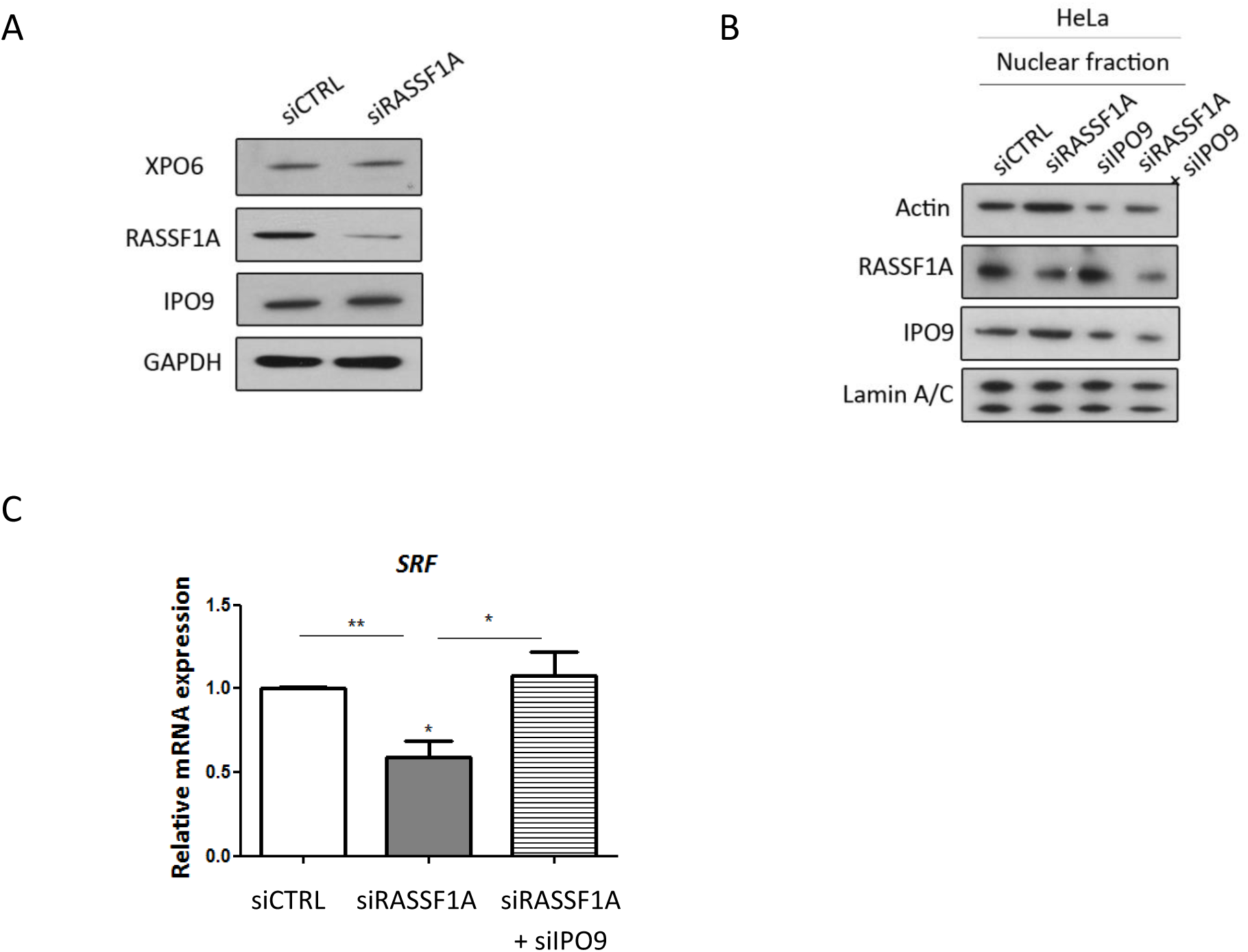
A. Western blot analysis of exportin 6 (XPO6) and importin 9 (IPO9) in the absence of RASSF1A. GAPDH was used as a loading control. B. Western blot analysis of actin protein levels following RASSF1A or RASSF1A/IPO9 knockdown in HeLa cells. C. SRF mRNA levels in siRASSF1A or siRASSF1A+siIPO9-transfected HeLa cells relative to GAPDH and normalised to control siRNA levels. Data represent the mean ± SEM of 3 replicates. *P <0.05, **P<0.01

**Supplementary Figure 4:**
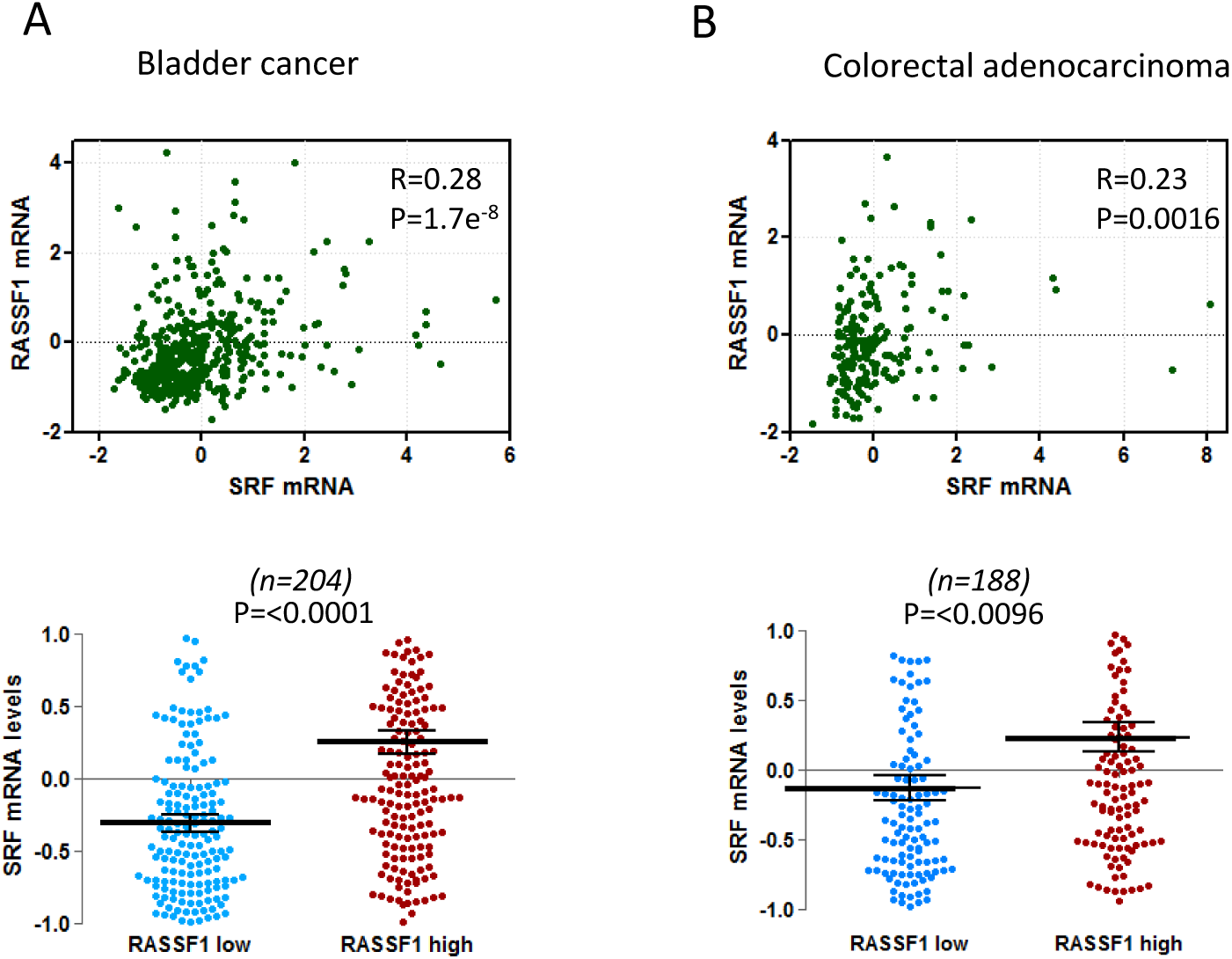
A. Upper: Scatter plot of RASSF1 mRNA to SRF mRNA levels in bladder cancer (TCGA, Cell 2017, n=404). Values are given in (RNA Seq V2 RSEM). Lower: Differences in SRF mRNA expression of samples that express low and high levels of RASSF1 mRNA. B. Upper: Scatter plot of RASSF1 mRNA to SRF mRNA levels in colorectal adenocarcinoma (TCGA, Nature 2012, n=195). Values are given in (RNA Seq V2 RSEM). Lower: Differences in SRF mRNA expression of samples that express low and high levels of RASSF1 mRNA.

